# Effects of green manure application on soil microbial communities and activities in the decontaminated sandy soil paddy field in Fukushima, Japan

**DOI:** 10.1101/2021.03.22.436377

**Authors:** Chol Gyu Lee, Yuko Mitsuda, Soh Sugihara, Taiichiro Ookawa, Haruo Tanaka

## Abstract

On March 11, 2011, Japan experienced an unprecedented earthquake off the Pacific coast of Tohoku, and suffered the direct and long-term effects of the earthquake and tsunami in the area. In Fukushima prefecture, agricultural land contaminated with radioactive Cesium from the Fukushima Daiichi Nuclear Power Plant. Therefore, surface soil were removed for deconamination, and low fertility sandy soil was covered. Organic matter input is necessary to increase soil organic matter and green manure application is an effective method to improve soil fertility in the paddy field. Soil microbes and enzyme activities are sensitively responded to organic matter addition, but their dynamics on the dressed field are not well investigated. In this study, we focused on changing the microbial community, diversity and enzyme activities along with the green manure decomposition process in the sandy soil dressed paddy field in Japan. The green manure of hairy vetch and oat were harvested and incorporated in May 2020 and their decomposition process as cellulose and hemicellulose contents were determined. Soil bacterial communities were analyzed using 16S amplicon sequencing. The green manure was rapidly decomposed within the first 13 days, and they did not remain 50 days after green manure incorporation. Soil microbial biomass carbon was higher in the M treatment after GM treatment, but was not significant between treatments after 50 days. Dehydrogenase and β-glucosidase activities changed during the harvesting period, but did not correlate with GM decomposition. Microbial diversity (OTU numbers and Shannon index) also changed with GM application, but they were not associated with GM decomposition. Soil prokaryotic communities and some bacteria (*Baciili* and *Chlorolfexi*) are significantly influenced by GM treatment. However, *Clostrida* was not affected by GM. Mixed green manure treatment showed significantly rapid hemicellulose decomposition than other treatments. In this process, *Anaerolineae* were negatively correlated with the decreasing of hemicellulose in this treatment. These results showed that GM treatment affected microbial communities, and their response was active during the decomposition process.

## Introduction

The Great East Japan Earthquake and tsunami on 11 March 2011, caused damage to the Fukushima Daiichi Nuclear Power Plant (NPP), resulting in serious radioactive pollution throughout Eastern Japan. The radioactive fallout extensively polluted agricultural lands, including paddy fields, with radioactive Cs (MEXT 2011). The Cs contaminated agricultural soil were removed for depth 10cm, and the sandy soils with low soil carbon and nitrogen content (total carbon is 8.29 g kg soil^-1^ and total nitrogen is 0.74 g kg soil^-1^) were covered as decontamination. The decline of soil organic carbon negatively impacts crop productivity and sustainability of agriculture (Lal 2004; Agegnehu et al. 2016). Organic amendments can provide available nutrients for plants, and the coupling of carbon and nutrient transformation during organic matter decomposition strongly interacts with plant nutrient uptake (Kaye and Hart, 1997). The application of organic materials to rice fields for yield increase has a long history in Asian countries. Recent studies have focused on re-considering traditional fertilization practices to enhance soil organic input by amendments of crop residues, green manure, and farmyard manure (Liu et al. 2009). The most useful organic matter in the paddy field is rice straw. However, the rice yields in this decontaminated paddy field in Fukushima are less than half the average yield in Japan, therefore, not enough rice straw can be applied to increase soil fertility. Livestock wastes are another important organic amendment; however, the stock rising was not restarted yet in this area. The application of green manure (GM) to paddy fields is considered a good management practice (Zhang et al. 2017). GM application has been reported to increase soil organic matter, and fertility, and nutrient retention, reducing the occurrence of plant disease and long- term green manure incorporation increases rice yields (Gao et al. 2013; Li et al. 2019). Soil microbes play an important role in maintaining soil fertility and productivity and drive most soil processes, e.g. decomposition of organic materials, nutrient availability and retention, and soil organic matter sequestration (Coleman et al., 2004). Soils with high fertility generally possess larger microbial biomass, higher enzyme activities, and better soil structure than those with low fertility (Fontaine et al. 2011; Lang et al. 2012) could provide a suitable environment for substrate utilization by microbes. However, microbial communities, abundance, and their activities that respond to plant residue decomposition in the sandy soil are still limited. For recovering SOM in the covered sandy paddy field after decontamination in Fukushima, it is essential to investigate microbial response to SOM decomposition. It was reported that the mixing of different species of GM can effectively improve soil fertility than a single GM application (Fageria et al., 2005; Tosti et al., 2014). The mixtures of plant residue at rates faster than expected from the average of the decomposition rates of the plant types component. This phenomenon is termed the "mixing effect" and the hypotheses proposed to explain it include physical, chemical, and biological mechanisms. The aim of this study is to investigate the effects of GM application on soil enzyme activity and microbial community during the GM decomposition process in sandy soil decontaminated paddy fields in Fukushima Prefecture.

## Material and Methods

### Sampling fields

The paddy field was located in Tomioka town, Fukushima Prefecture, Japan (37°20’N, 140°60’E). The fields are about 12km away from Fukushima 1st NPP. The soil was sandy loam soil (Fluvaquents) in the top 0.5 m, with 78.7 % sand, 17.7 % silt and 3.6 % clay. Soil physicochemical properties in the field were shown in Table 1. Two species of green manures were used, i.e., oat (A: *Avena strigosa* cv. Hayoats) and hairy vetch (V: *Vicia villosa* cv. Fujiemon). These species were sown as pure crops at an ordinary sowing rate (4 kg of hairy vetch and 15 kg oat each in 10a^-1^) and as a mixture with the same quantity (M: 4 kg of hairy vetch and 15 kg of oat 10a^-1^). Non-green manure treatment plots (control: C) were included in the experiments. The experimental design was a randomized complete block with three replicates. Each plot size was 225 m^2^ (15 m×15m). GM was cultivated from 1 November 2018 to 5 May 2019.

**Table 1.**
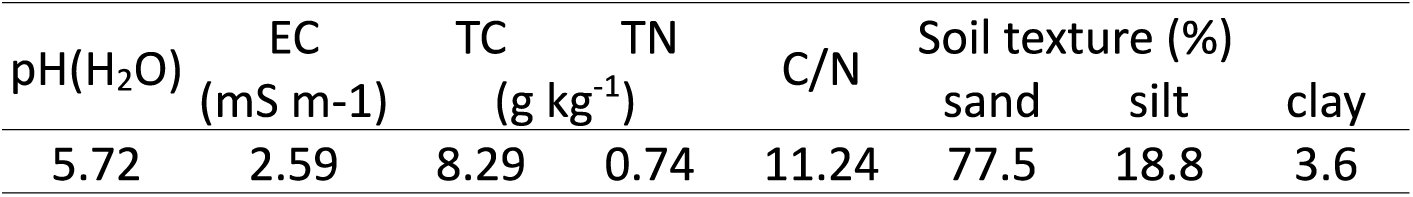
Soli physicochemical properties of the paddy field

### Litterbag experiments

Each green manure was harvested from randomly selected 2 points (0.5m × 0.5m) in each treatment on 5 May. The yields and chemical properties of each GM were shown in Table 1. The GM dried at 70 °C and powdered. The litterbags (1mm mesh size) were filled with 30g of air-dried paddy field soil mixed with each GM. The amount of each GM input was equal to the amount of input carbon rate in the field (0.05 % of O and V treatment and 0.11% of M treatment) (Table 2). A total of 72 litterbags (4 treatment × 6 sampling times × 3 replicants) were prepared. All litterbags were incorporated in each control plot of the paddy field on 30 May 2020. Within each plot, 24 litterbags (4 treatments and 6 were incorporated into the soil by burying them at 15cm depth. Each litterbag from each plot was removed chronologically from 12 June (13 days), 3 July (34 days), 30 July (50 days), 21 August (72 days), and 4 October (116 days). One gram of soil in the litterbags were transported to the laboratory using an icebox and processed immediately after their removal from the field.

**Table 2.** Yields and chemical properties of each GM

|  | Yields | TC | TN | CN ratio |
| --- | --- | --- | --- | --- |
|  | t/ha | t/ha | t/ha |  |
| V | 1.24 | 0.53 | 0.045 | 11.8 |
| A | 1.31 | 0.61 | 0.01 | 61.0 |
| M | 2.55 | 0.9 | 0.03 | 30.0 |

### Soil chemical properties

Total carbon and nitrogen contents were measured with a NC analyzer (SUMIGRASH NC-80, Sumitomo Chemical Co. Ldt., Tokyo, Japan). Cellulose and hemicellulose content were determined with colormetric anthrone-sulfuric acid method (Koehler 1952) after hydorolysis of component sugars by Oades et al. (1970). The cellulose and hemicellulose content of the control at that time was subtracted from each treatment section to obtain the cellulose and hemicellulose content at that time. The content at the time of treatment was set to 100, and the degradation rate was determined.

### Soil biological properties

Soil enzyme activity was determined as below. Dehydrogenase activity was determined with the reduction of iodonitrotetrazolium chloride (INT) by Von Mersi and Schinner (1991). β-glucosidase activities were assayed on the basis of p-Nitrophenyl-β- D-glucopyranoside (PNG) hydrolysis after cleavage of enzyme-specific synthetic substrates by Hayano (1973). Microbial biomass carbon and nitrogen were determined by chloroform fumigation-extraction method with 0.5 M K_2_SO_4_ at 1:4 soil to extraction ratio (Moore et al. 2000)

### DNA extraction and microbial community analysis

DNA was extracted from 0.5 g of soil using the ISOIL for bead beating kit (Nippon Gene Co., Ltd., Tokyo, Japan), according to the manufacturer’s instructions. DNA quantification and integrity were measured using a NanoDrop spectrophotometer (Thermo Fisher Scientific, Waltham, MA, USA) and gel visualization (0.8% agarose in Tris/acetic acid/ethylenediaminetetraacetic acid buffer), respectively. The V4 region of the 16S rRNA gene of each sample was amplified by PCR using the bacterial and archaeal universal primers 515F (5′-GTGCCAGCMGCCGCGGTAA-3′) and 806R (5′-GGAC- TACVSGGGTATCTAA-3′) (Caporaso *et al*. 2011). A library was prepared by adaptor ligation with the PCR primer pairs using the TruSeq Nano DNA Library Prep Kit (Illumina, Inc., San Diego, CA, USA). When two or more bands were detected using 1.5%-agarose gel electrophoresis, PCR products of approximately 300 bp in length were excised from the gel, non-specific amplicons were removed, and the products were purified using a MonoFas DNA purification kit for prokaryotes (GL Sciences, Inc., Tokyo, Japan). Each PCR amplicon was cleaned twice to remove the primers and short DNA fragments using the Agencourt AMPure XP system (Beckman Coulter, Inc., Brea, CA, USA) and quantified using a Qubit Fluorometer (Invitrogen Corporation, Carlsbad, CA, USA). The PCR products were adjusted to equimolar concentrations and subjected to unidirectional pyrosequencing, which was performed by Bioengineering Lab. Co., Ltd. (Kanagawa, Japan) using a MiSeq instrument (Illumina, Inc.). Overall, 3,521,651 sequences were obtained from the 72 samples (Supplemental Table 1). Sequencing data were deposited in the DNA Database of Japan Sequence Read Archive under the accession number DRA006673.

### Statistical analysis

Illumina sequence data were sorted based on unique barcodes and quality-controlled using the Quantitative Insights Into Microbial Ecology Qiime2 (version 2017.8, https://docs.qiime2.org/2017.8/) with plugins demux (https://github.com/qiime2/q2-demux), dada2 (Callahan et al., 2016) and feature-table (McDonald et al., 2012). Alpha and beta diversity analyses were performed by using plugins alignment (Katoh and Standley, 2013), phylogeny (Price et al., 2010), diversity (https://github.com/qiime2/q2-diversity), and emperor (Vazquez-Baeza et al., 2013). A pre-trained Naive Bayes classifier based on the Greengenes 13_8 99% OTUs database (http://greengenes. secondgenome.com/), which has been trimmed to include the v4 re- gion of 16S rRNA gene, bound by the 515F/806R primer pair, was applied to paired-end sequence reads to generate taxonomy tables. Taxonomic and compositional analyses were conducted by using plu- gins feature-classifier (https://github.com/qiime2/q2-feature- classifier), taxa (https://github.com/qiime2/q2-taxa) and composition (Mandal et al., 2015). The significant difference among each treatment was analyzed by Tukey-Kramer HSD method using R. Beta diversity was measured according to Bray–Curtis distances which were calculated by R, and displayed using Principal Coordinate Analysis (PCoA). The significance of grouping in the PCoA plot was tested by analysis of similarity (ANOSIM) in R with 999 permutations. The interacted factor between cellulose, hemicellulose, and carbon decomposition and related microbes were analyzed using Spearman’s rank correlation. The correlation coefficients (Rho value) less than -0.4 were defined as negatively correlated microbes and more than 0.4 were defined as positively correlated microbes.

## Results

### The dynamics of GM content during the harvesting period

GM, which mainly consists of cellulose and hemicellulose, was mainly degraded in 31 days after GM treatment. At the 13 days, 56-72% of cellulose was decomposed, and the degradation rate did not differ among the treatments (Fig. 1). After 31 days, 5 to 14% of cellulose remained in all treatments and they remained unchanged through the harvesting period. Hemicellulose contents also decreased significantly over 13 days, but they differed among treatments: M treatment, a mixture of V and A treatment, decreased significantly (99%) compared to the other treatments (50% in V and 23% in A). Thereafter, the residues remained below 10% until day 116. In the V treatment, the rate of residues reached 33% on day 72, but they did not significantly differ from the other treatments. Total carbon and total nitrogen contents were gradually decreased in all treatments and the highest in M on 34 days after GM treatment (Figure 1). After 50 days, TC and TN contents were similar among each GM treatment. Therefore, GM might be completely decomposed until this time. Total carbon content in each field was also changed during the harvesting stage, though they were not different between the treatments (Supplemental figure 1). After the rice harvesting, soil carbon and nitrogen content, cellulose, and hemicellulose were not different among the treatment (Table 3).

**Figure 1.**
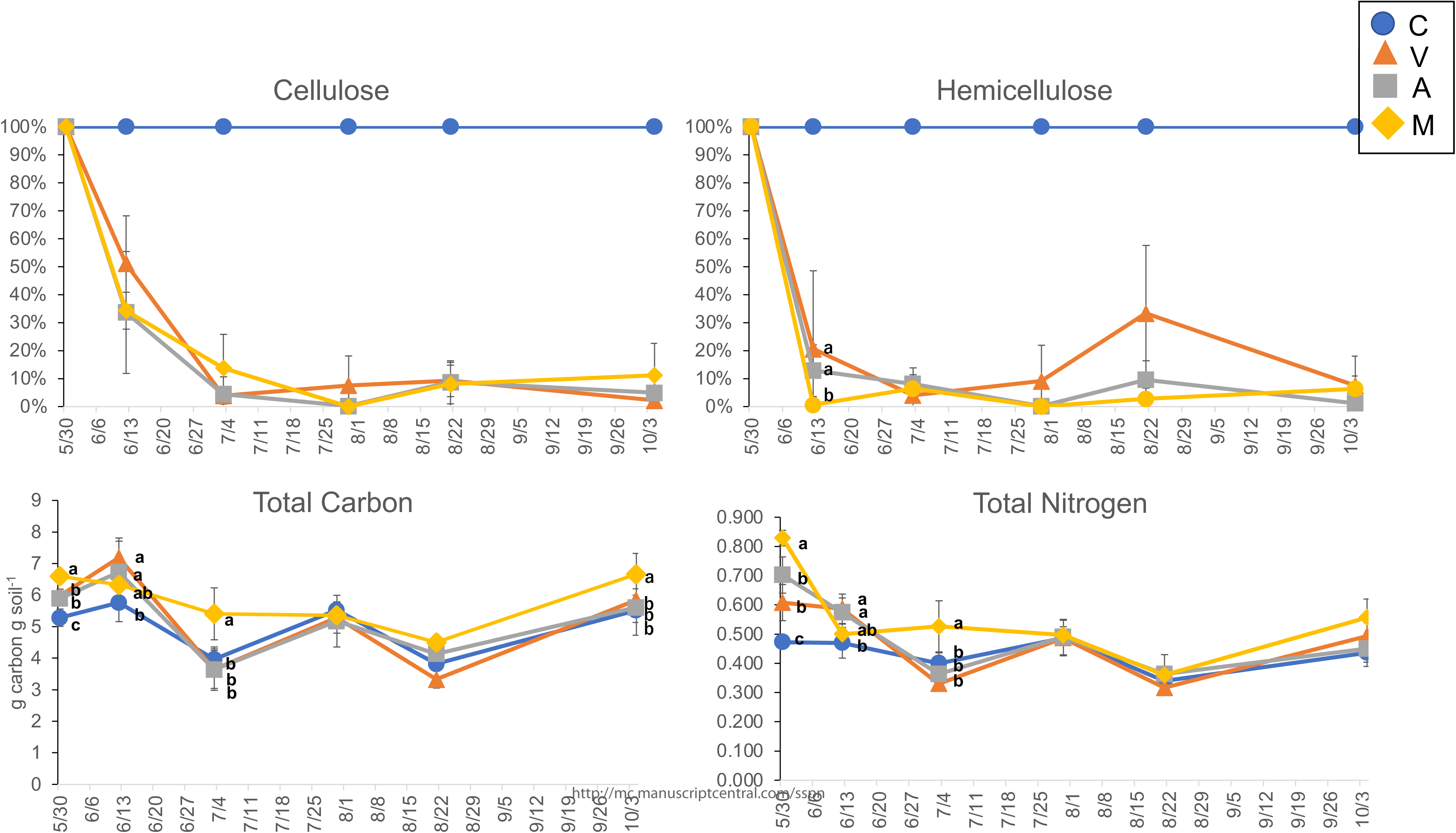
Dynamics of cellulose, hemicellulose, total cabon and nitrogen contents in each treatment during the harvesting period. Error bars indicate standard deviations. Statistically significant treatments in each sampling days are indicated by alphabetic labels (Tukey HSD analysis, p < 0.05).

**Table 3.**
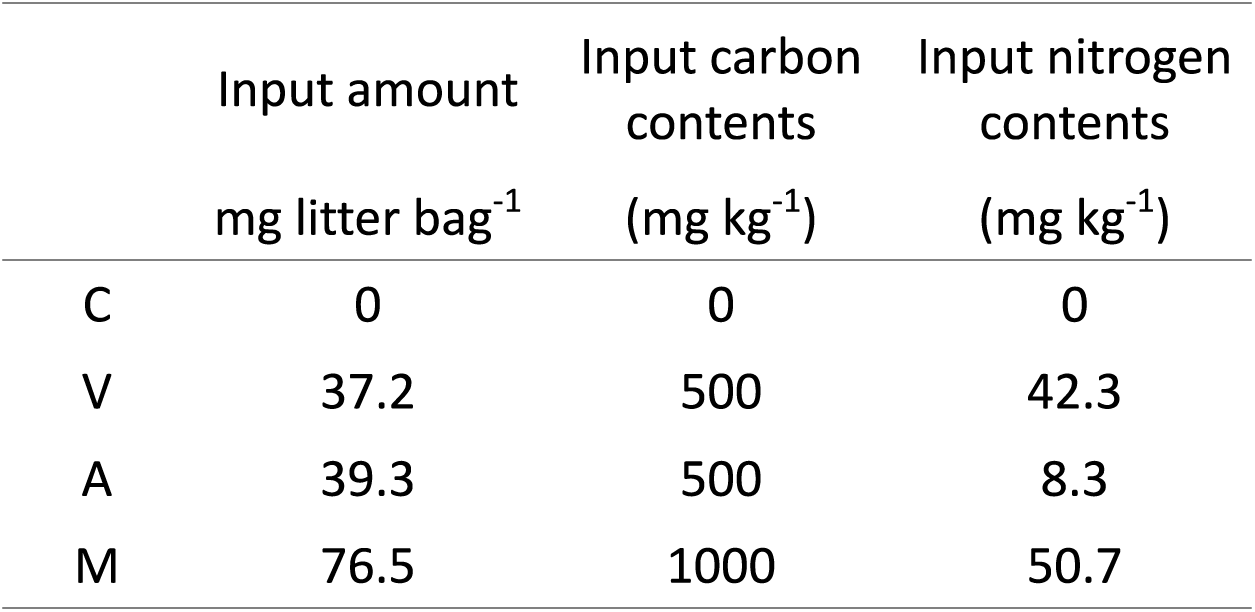
Inputs of GM in the litter bag

|  | Input amount | Input carbon | Input nitrogen |
| --- | --- | --- | --- |
|  | mg litter bag<sup>-1</sup> | contents | contents |
|  |  | (mg kg<sup>-1</sup>) | (mg kg<sup>-1</sup>) |
| C | 0 | 0 | 0 |
| V | 37.2 | 500 | 42.3 |
| A | 39.3 | 500 | 8.3 |
| M | 76.5 | 1000 | 50.7 |

### Dynamics of microbial biomass, community structure, diversity, and enzyme activities response to GM application

Soil microbial biomass carbon was significantly increased in M treatment followed by A, V, and C treatment on 13 and 34 days after GM treatment (Figure 2). They significantly decreased and were not significantly different among the treatments after 50 days. Dehydrogenase activities were increased 34 days after treatment, and they in A treatment showed the highest activities than other treatments. After 50days, the activities decreased and were not significantly different among the treatment. Beta- glucosidase activities were drastically changed during the harvesting stage; however, they were not significantly different among the treatments. Soil microbial diversity, OTU numbers and Shannon index, did not differ among treatments on days 13 and 34. They were the highest in M treatment compared to other treatments on 50 days and then decreased. On day 72 and 116, they in V treatment were higher than A treatment and M treatment, respectively.

**Figure 2.**
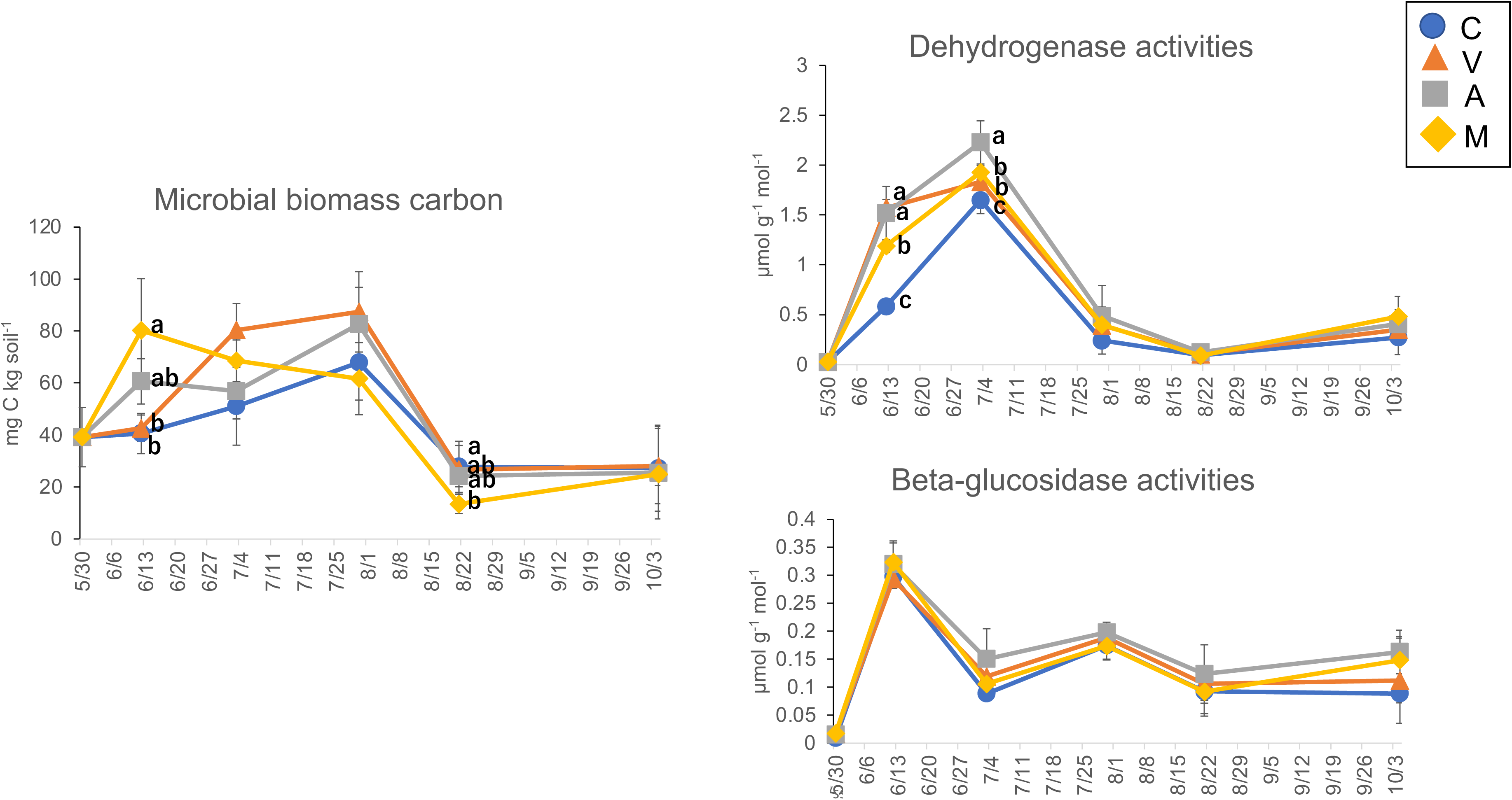
Dynamics of microbial biomass, and each enzyme acticity during the harvesting period. Error bars indicate standard deviations. Statistically significant treatments in each sampling days are indicated by alphabetic labels (Tukey HSD analysis, p < 0.05).

**Figure 3.**
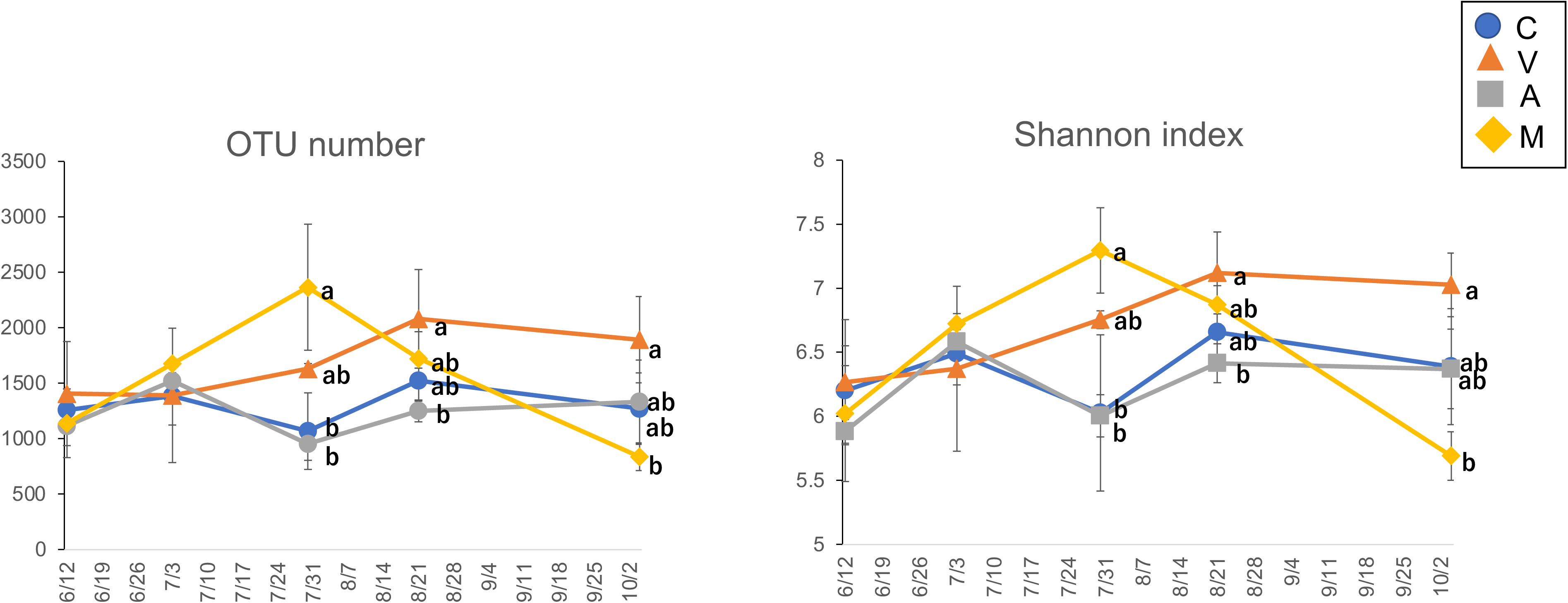
Dynamics of microbial diversity and richness during the harvesting period. Error bars indicate standard deviations. Error bars Statistically significant treatments in each sampling days are indicated by alphabetic labels (Tukey HSD analysis, p < 0.05).

The main prokaryotic phyla in the field were Firmicutes and Proteobacteria (Figure 4). They were occupied more than 50% in all treatments and sampling times. In class level, Bacilli were the most dominant bacteria in all treatments followed by Clostridia, Alphaproteobacteria, and unidentified Proteobacteria (Figure 5). The relative abundance of *Bacilli* was decreased on day 31 compared with that on 13 days, then was again high in the C and A treatments at day 50. Chloroflexi was higher in the M treatment than in the V treatment at day 50; the relative abundance of *Clostridia* did not differ among treatments (Figure 6). Soil prokaryotic communities were affected by GM treatment (anosim p < 0.05). PCoA analysis based on Bray-crutis analysis showed that prokaryotic communities were roughly clustered by sampling times (Figure 7).

**Figure 4.**
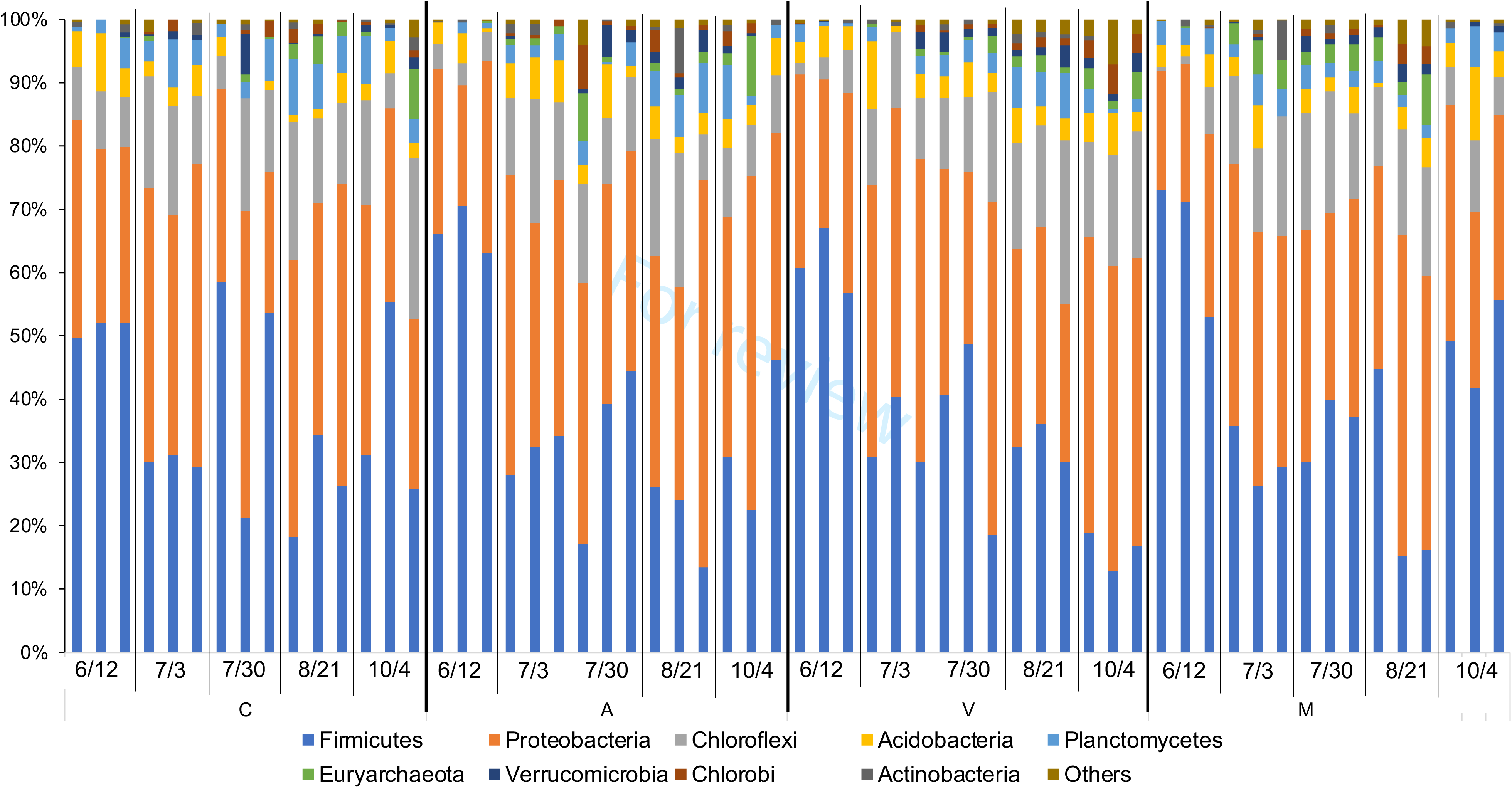
Changing the relative abundance of prokayoric communities during the harvesting period in genus level. The date indicated the sampling days and the alphabet at the bottom indicated each treatment.

**Figure 5.**
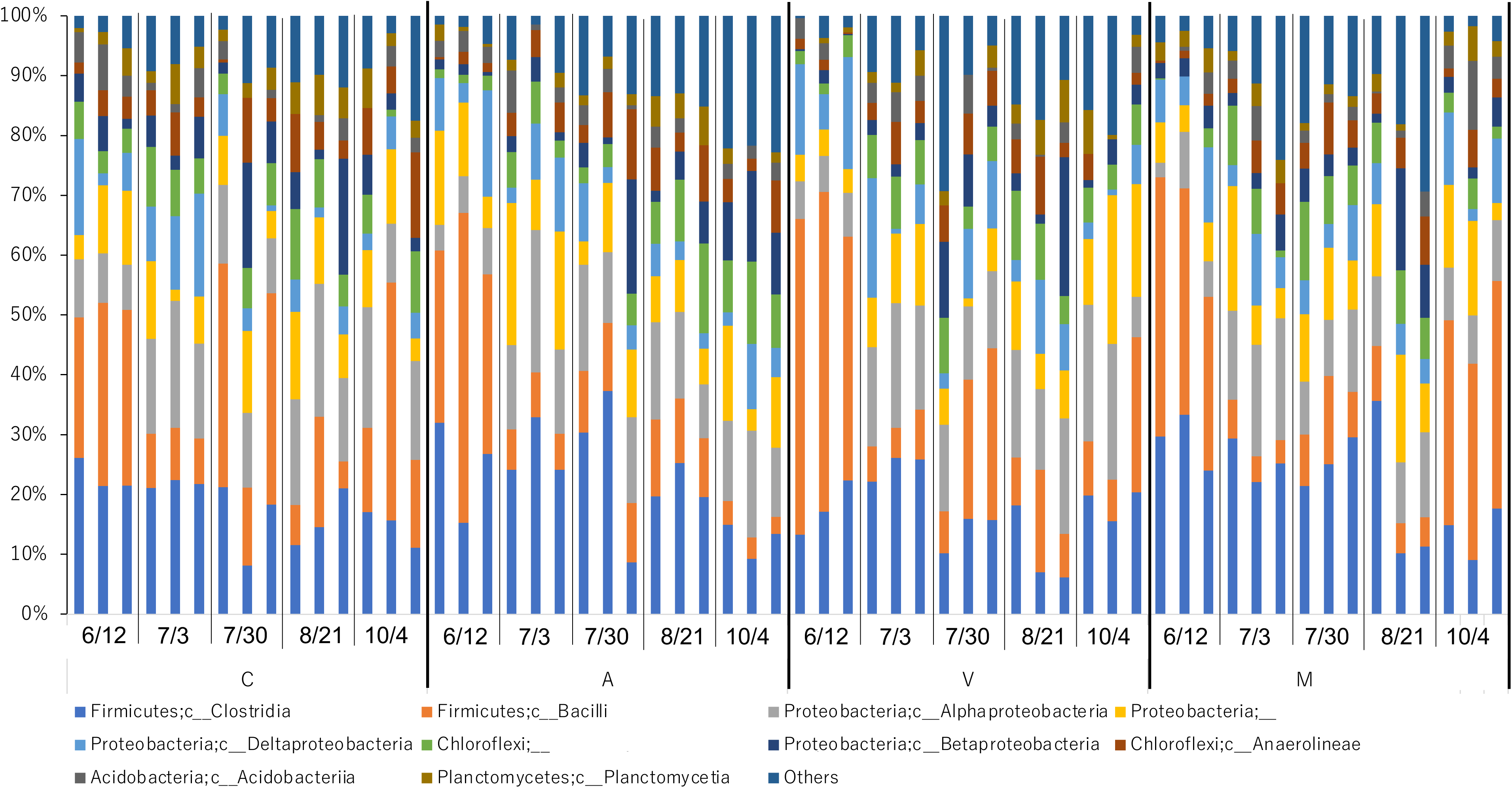
Changing the relative abundance of prokayoric communities during the harvesting period in order level. The date indicated the sampling days and the alphabet at the bottom indicated each treatment.

**Figure 6.**
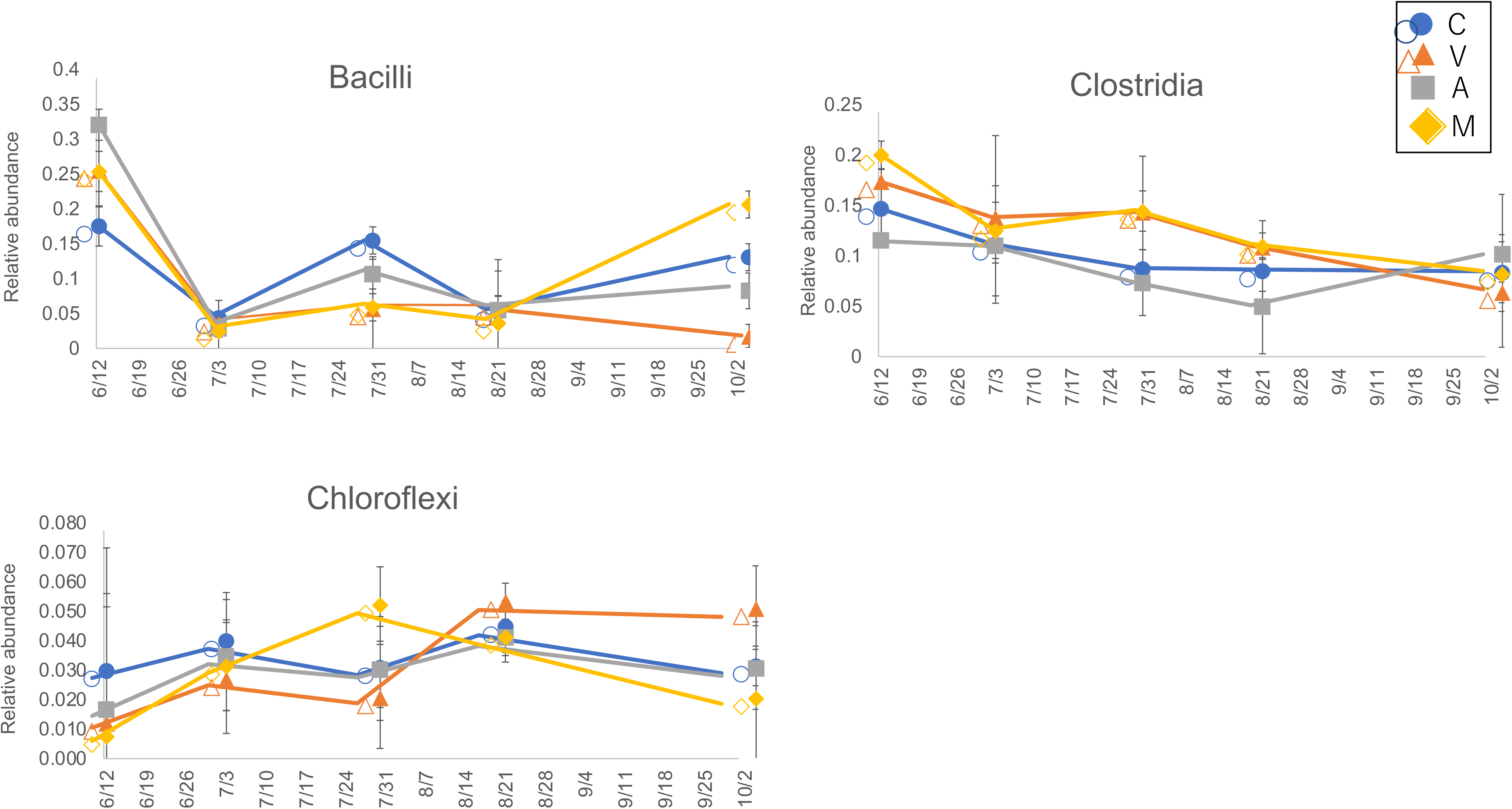
Dynamics of each microbes during the harvesting period. Error bars indicate standard deviations. Statistically significant treatments in each sampling days are indicated by alphabetic labels (Tukey HSD analysis, p < 0.05).

**Figure 7.**
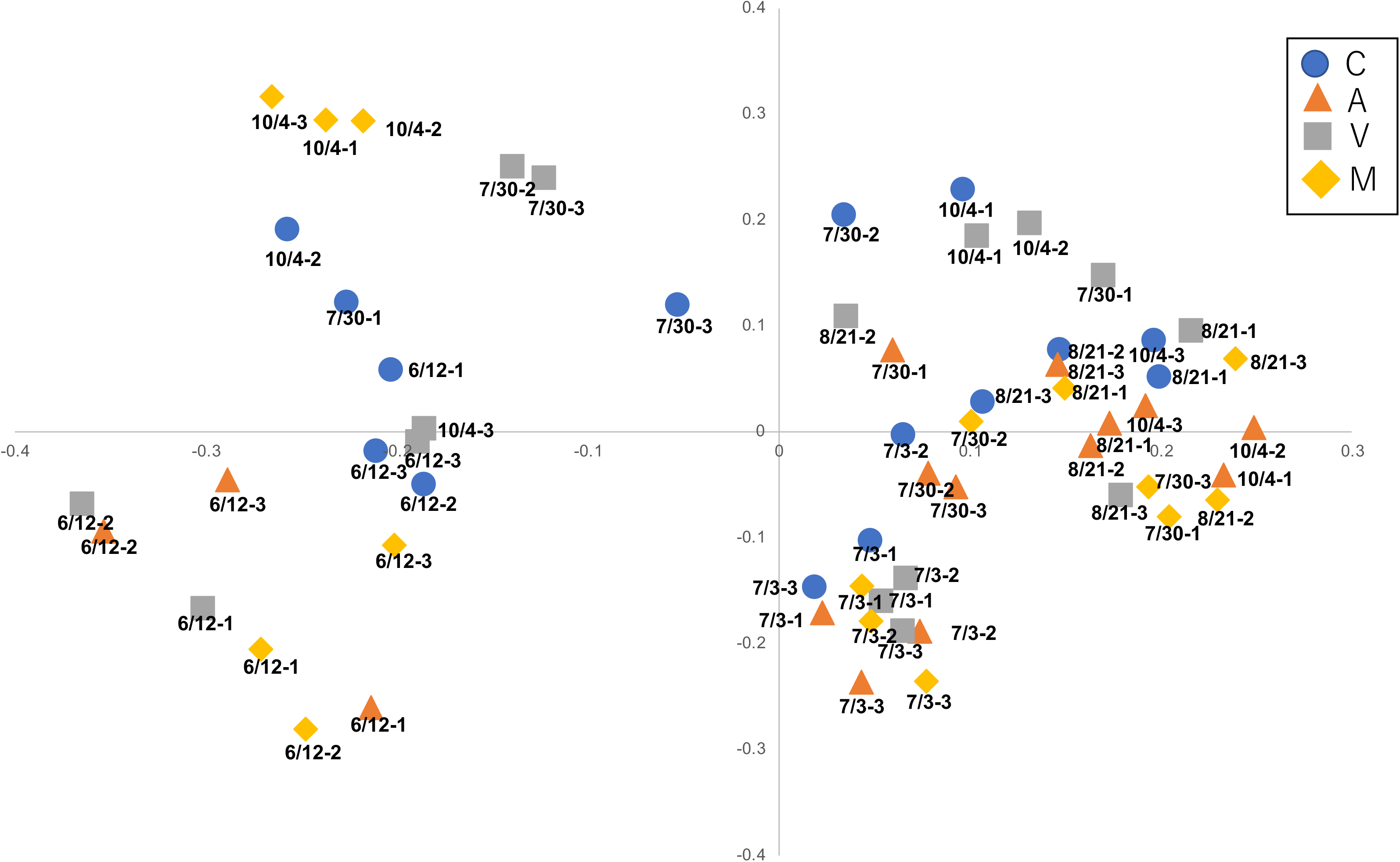
Beta diversity: principal coordinate analysis (PCoA) of prokaryotic community structure based on Buray–Curtis distances for each treatments during the harvesting period. Sample names were assigned x/y-z in which x, y and z indicate the date of sampling and replication number, respectively (ex, 6/12-1 was sampled at 12 June and the replication 1).

Prokaryotes sampled in 13 and 31 days were clustered by sampling times. But their communities were not clustered in sampling time after 50 days. We analyzed the microbes correlated between total carbon, hemicellulose, and cellulose contents (Table 4). *Lachnospiraceae* and *Clostridiales* belonged to Clostridia positively correlated to cellulose and hemicellulose content, and *Bacillus* correlated to total carbon content.

**Table 4.**
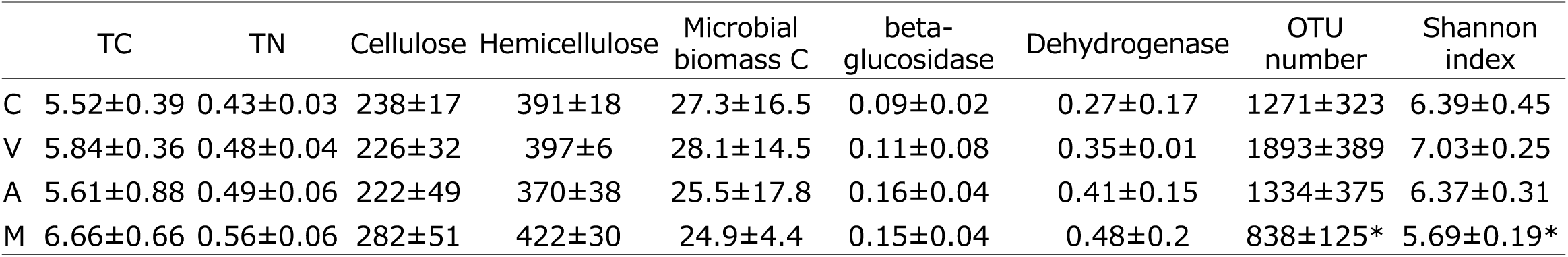
Soli chemical and microbial properties after ther harvesting stage

|  | TC | TN | Cellulose | Hemicellulose | Microbial<br>biomass C | beta-glucosidase | Dehydrogenase | OTU<br>number | Shannon<br>index |
| --- | --- | --- | --- | --- | --- | --- | --- | --- | --- |
| C | 5.52±0.39 | 0.43±0.03 | 238±17 | 391±18 | 27.3±16.5 | 0.09±0.02 | 0.27±0.17 | 1271±323 | 6.39±0.45 |
| V | 5.84±0.36 | 0.48±0.04 | 226±32 | 397±6 | 28.1±14.5 | 0.11±0.08 | 0.35±0.01 | 1893±389 | 7.03±0.25 |
| A | 5.61±0.88 | 0.49±0.06 | 222±49 | 370±38 | 25.5±17.8 | 0.16±0.04 | 0.41±0.15 | 1334±375 | 6.37±0.31 |
| M | 6.66±0.66 | 0.56±0.06 | 282±51 | 422±30 | 24.9±4.4 | 0.15±0.04 | 0.48±0.2 | 838±125* | 5.69±0.19* |

Cellulose contents were negatively correlated (*ρ* < -0.4) *Anaerolineae SJA15* and unidentified Chloflexi belonged to Chloflexi, and *Rhizobiales* belonged to Alphaproteobacteria. Hemicellulose contents were negatively correlated to *Anaerolineae SJA15* and unidentified Chloflexi belonged to Chloflexi, unidentified betaproteobacteria, *Pedosphaera* belonged to Verrcomicrobia, Chrolobi, and *Methanomicrobia* belonged to Euryarchaeota. Total carbon was negatively correlated with *Ktedonobacteria* and unidentified Chloroflexi belonged to Chloroflexi, *Rhizobiales* belonged to Alphaproteobacteria, and *Chlorobi BSV26*. We compared the correlation between hemicellulose content and each microbe among the GM treatments (Table 5). *Bacillu*s were positively correlated to hemicellulose content in all treatments, and *Lachnospiraceae* belonging to Clostridia positively correlated with E and M treatment. On the other hand, Anaerolineae were negatively correlated in A and E treatment, respectively, but not in M treatment. Some methanogen (*Methenomicrobia* and *Methanogulaceae*) were negatively correlated with hemicellulose content only in M treatment.

**Table 5.**
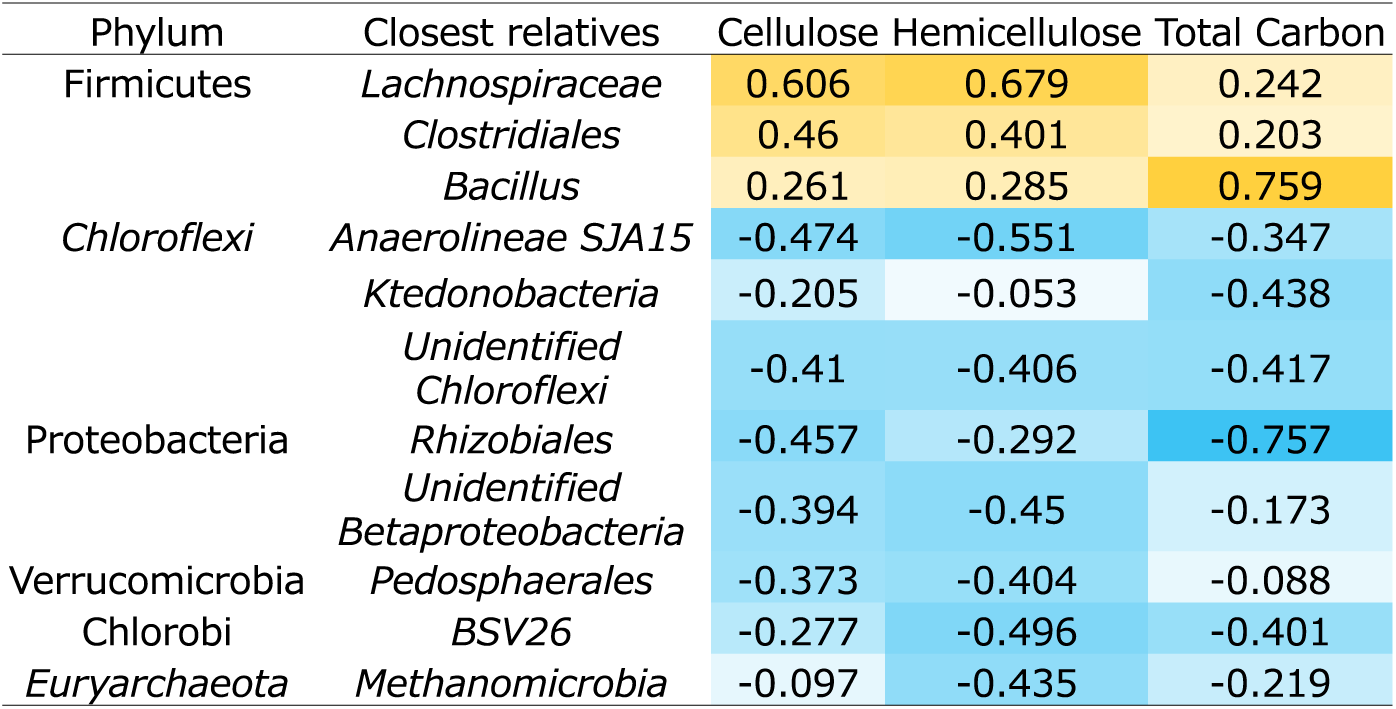
Correlations between each microbes and cellulose, hemicellulose, and soil carbon content

**Table 6.**
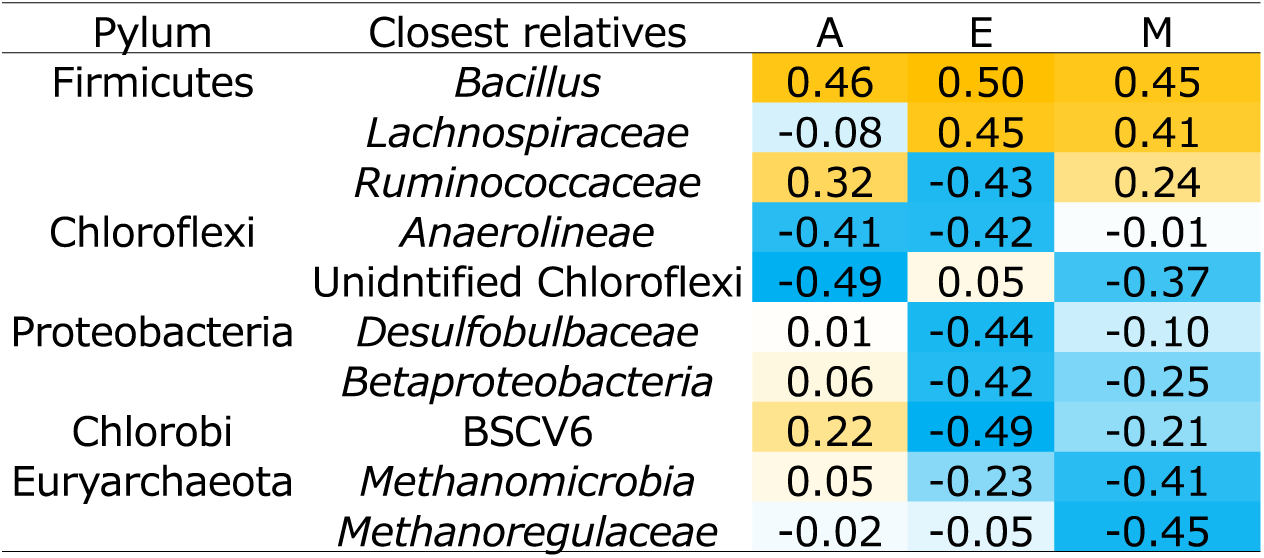
Correlations between each microbes and, hemicellulose contents in each treatment

## Discussion

### Microbial communities and activities during GM decomposition process

GM is used as a useful organic matter in the paddy field and it increases soil organic carbon and nitrogen (Lee et al. 2010; Hwang et al. 2015). Gao et al. (2016) reported that about 70% of wheat residue was mineralized under the anaerobic condition as well as aerobic condition in a Cambisol after 12-month in a field experiment. Zech et al. (1997) concluded that 30-45 % of plant residue carbon was accumulated in the soil after 1 year treated with various SOM conditions. In Japanese field, 67 to 79 % of carbon from rice residues was decomposed during one year (Shiga et al. 1985). On the other hand, in our fields, cellulose and hemicellulose were decomposed within 50 days and soil carbon was not accumulated after the rice harvesting period. The aboveground biomass of green manure (barley and hairy vetch) in past related studies ranges from 3.66 to 11 t ha^-1^ and is incorporated into the field (Tosti et al. 2014; Jeon et al. 2008, Hwang et al. 2015). However, in our field, only 1.24-2.55 t ha^-1^ of GM was harvested due to the cold climate of Fukushima and low soil fertility of the covered sandy soil (Table 1). Another possibility of the rapid GM decomposition is the sandy textured soil property. Most sandy soils have low soil organic matter content and show low water holding capacity and high permeability due to their coarse texture (Rutkowska and Pikuła, 2013; Šimanský and Bajčan, 2014). Some studies showed that organic amendments generally decompose rapidly in sandy soils due to high temperatures and aeration (Glaser et al., 2002). Xu et al. (2019) showed that mineralization of residue carbon after 360 days of incubation was higher in low fertility soils than in high fertility soils. Low fertility soils have a higher C/N ratio than high fertility soils. In this case, soil microorganisms that are deficient in nitrogen nutrients will use nitrogen derived from the residue pool, which is more easily decomposed, and will preferentially decompose plant residues to meet their nitrogen requirements. These two factors might contribute to the rapid degradation of GM.

Dehydrogenase is an intracellular enzyme that participates in oxidative phosphorylation in microorganisms and basically depends on the metabolic state of the soil microbes (Tabatabai et al. 1994; Insam, 2001). β-glucosidase is an extracellular enzyme that contributes to the degradation of cellulose and other β-1,4 glucans to glucose and is considered to be one of the proximate agents of organic matter decomposition (Sinsabaugh et al., 2008). Soil organic matter is the substrate for these soil enzymes, therefore, dehydrogenase and beta-glucosidase are positively correlated with soil organic matter (Bhattacharyya et al. 2012). However, in our study, each enzyme activity was not correlated with soil carbon, cellulose, and hemicellulose contents. Moreover, no microbes were correlated with each enzyme activity. The soil microbial biomass carbon in this study (13.4 to 87.4 mg C kg soil^-1^) was much lower than that in the previous study (more than 200 mg C kg soil^-1^) (Lu et al. 2002; Bhattacharyya et al. 2012; Zheng et al. 2016). Because the amount of green manure input was low, the small amount of enzymes may be secreted, and no difference was observed between the treatment. Moreover, β-glucosidase is an extracellular enzyme; therefore, it was leached from the outside of the litter bag in the flooded paddy field and contaminated each treatment.

*Clostridia*, organic matter degradation bacteria under anaerobic conditions, is known to increase during the degradation process under flooded conditions (Weber and Conrad 2001; Shrestha *et al*. 2011). Lee *et al*. (2017) showed that dried rice callus cells, a model of easily degradable plant residues, are mainly degraded by *Clostridia* under anaerobic conditions. In contrast, the abundance *Clostridia* did not change during the harvest period and was positively correlated with cellulose and hemicellulose content. *Bacillus*, the dominant aerobic bacteria in the field, was also significantly reduced during the harvest period. Soil condition might be anaerobic and it is favorable for *Clostridia* increasing. *Clostridia* and *Bacilli* are known for rapidly increasing bacteria response to organic matter addition (Bao *et al*. 2019). Therefore, it was suggested that they might be involved in the fast decomposition of GM and then they dominated.

Thereafter, they could not grow because of low substrates. Compared with their decreasing, bacteria belonging to *Chloroflexi, Rhizobiales, Betaproteobacteria,* and *Chlorobi* BSV26 negatively correlated to soil carbon and plant cell wall content. They are well-known oligotrophic bacteria (Fierer *et al*. 2007; Tian et al. 2015). They can grow in the low nutrient soil as oligotrophs, nitrogen-fixing, sulfate-reduction, and photosynthesis. In this study, there was no effect of GM treatment on carbon accumulation in soil. On the other hand, we revealed that the soil microbial community was affected by GM treatment. The microbial community associated with carbon from GM accumulation was also increased. In the future, the microbes involved in carbon sequestration and link it to the implementation of soil improvement and rice production in the covered field.

### Mixing effect on GM decomposition and microbial properties

The types of GM also affected their decomposition rate. In general, plant residue decomposition is correlated with C/N ratio of each plant (). The combination of barley and hairy vetch optimizes the C/N ratio, which can favor the mineralization of organic substrates in soil (USDA, 2011). In this study, C/N ratio of V was 12 and that of E was 61, therefore, we hypothesized the decomposition rate was V > M > E. Cellulose decomposition rates were not different among the treatment, while M treatment showed the highest hemicellulose decomposition rates compared with other treatments.

Chapman and Koch et al. (2007) showed mixtures of plant litter decomposed up to 50% faster than expected decomposition speed and single litter types. Cuchietti et al (2014) explained that the fast-slow mixture decomposition rate is greater than expected due to the fast decomposing GM. In chemical aspects, the carbon limitation for decomposing fast decomposing plant residue and the slow decomposing plant residue it contains high CN ratio, will supply the carbon for carbon decomposition, therefore, decomposition will be accelerated. They indicated that fast-decomposing species would transfer nutrients to slow decomposing species, thereby increasing the decomposition rate of the slow decomposition species. Microbial communities also influence the mixing effect.

Our results showed that microbes affected by hemicellulose decomposition in single and mixed treatment showed differently. Methanogens negatively correlated with hemicellulose content only in M treatment. They are faculty anaerobic, use H_2_/CO_2_ and formate as a substrate for methanogenesis, and acetate is required for growth. The microbes involved in the mixing effect are not well studied. Our study provides some evidence that can shed light on some mechanisms of in the sandy soil environemnt, but on microbial and to the mechanisms proposed is critical.

## Acknolwgement

We thank Mr. Nobiru Watanabe for his supporing for management the paddy field. We alslo thank the members of Soil Science laboratory and crop production laboratory of Tokyo University of Agriculture and Technology for helping green manure sowing incorporating. This work was supported by the Fukushima Innovation Coast Framework Promotion Project and in part by the the Japan Society for Promotion of Science (JSPS) Grant-in-Aid for Scientific Research Grant Number 20H05583 (Post-Koch Ecology) and No.20H03113 (Scientific Research B).

---

**Figure S1.**
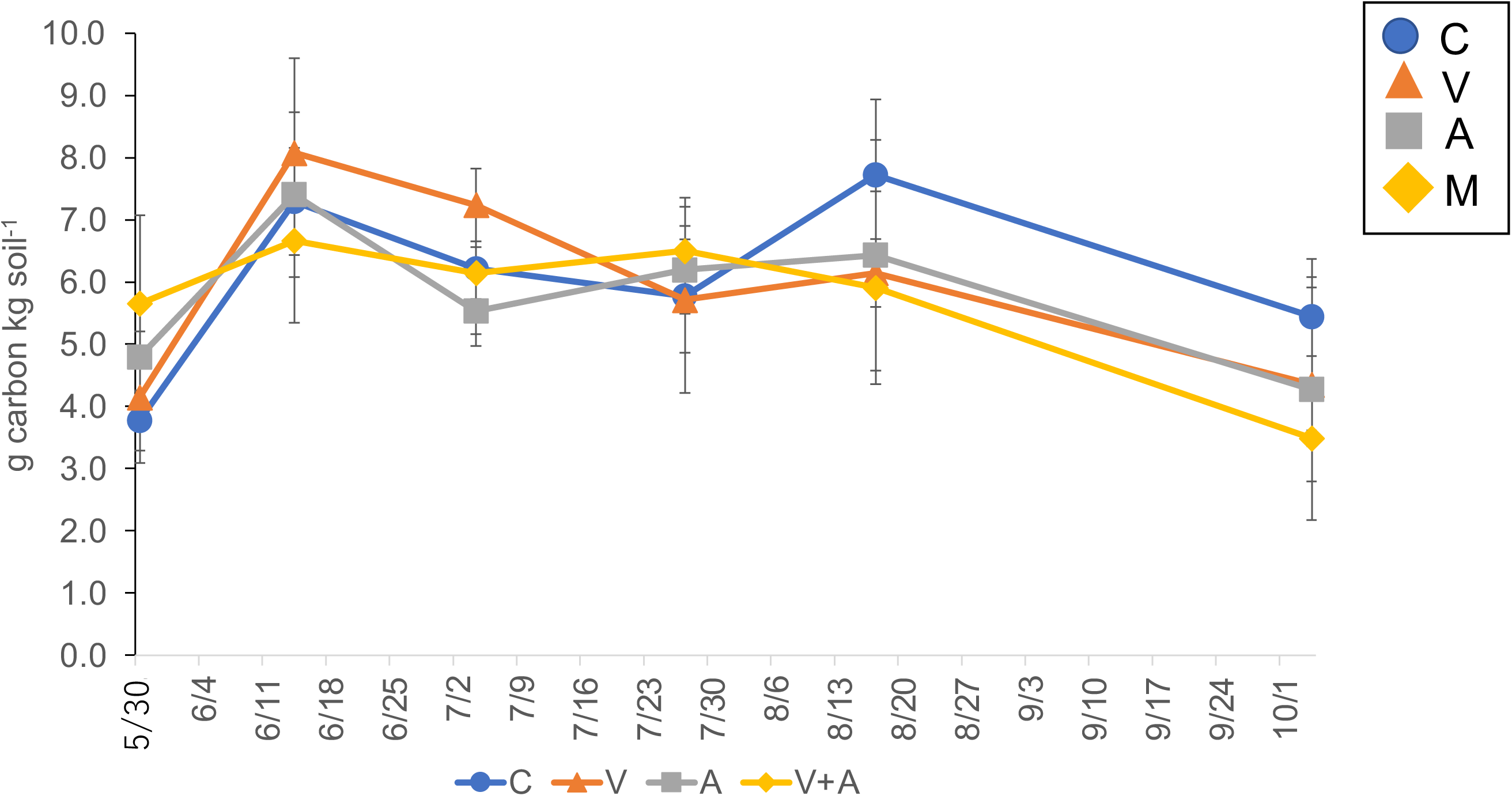
Changing the soil carbon content in the paddy field.

